# Automatic classification of constitutive and non-constitutive metabolites with gcProfileMakeR

**DOI:** 10.1101/2020.02.24.963058

**Authors:** Fernando Pérez-Sanz, Victoria Ruiz-Hernández, Marta Isabel Terry, Sara Arce-Gallego, Julia Weiss, Pedro J Navarro, Marcos Egea-Cortines

**Author notes:** VR-H and FP-S Contributed equally to this work. List of author contributions: V.R-H, S.A.G, P.J.N, F.P.S and M.E-C conceived the software; F.P-S, V.R-H and M.E-C conceived the original screening and research plans; F.P-S, S.A-G, and P.J.N coded the application; V.R-H, M.I.T, S.A-G, J.W, and M.E-C performed the experiments and analysed the data; V.R-H, M.I.T, J.W and M.E-C. wrote the manuscript. All authors corrected the manuscript. JW, ME-C and PJN wrote the grant applications. M.E-C. agrees to serve as the author responsible for contact and ensures communication. F.P-S and V.R-H contributed equally to this work. VRH Department of Biosciences, University Salzburg, 5020 Salzburg, Austria. SAG Vall d’Hebron Institute of Oncology, 08035 Barcelona, Spain.

## Abstract

Data analysis in non-targeted metabolomics is extremely time consuming. Genetic factors and environmental cues affect the composition and quantity of present metabolites i.e. the constitutive and non-constitutive metabolites. We developed gcProfileMakeR, an R package that uses standard output files from GC-MS for automatic data analysis using CAS numbers. gcProfileMakeR produces three outputs: a core or constitutive metabolome, a second list of compounds with high quality matches that is non-constitutive and a third set of compounds with low quality matching to MS libraries. As a proof of concept, we defined the floral scent emission of *Antirrhinum majus* using wild type plants, the floral identity mutants *deficiens* and *compacta* as well as RNAi lines of *AmLHY*. Loss of petal identity was accompanied by appearance of aldehydes typical of green leaf volatile profiles. Decreased levels of *AmLHY* caused a major increase in volatile complexity, and activated the synthesis of benzyl acetate, absent in WT. Furthermore, some volatiles emitted in a gated fashion in WT such as methyl 3,5-dimethoxybezoate or linalool became constitutive. Using sixteen volatiles of the constitutive profile, all genotypes were classified by Machine Learning with 0% error. gcProfileMakeR may thus help define core and pan-metabolomes. It enhances the quality of data reported in metabolomic profiles as text outputs rely on CAS numbers. This is especially important for FAIR data implementation.

**One sentence summary:** gcProfileMakeR allows the automatic annotation of the core metabolome and non-constitutive metabolites, increasing speed and accuracy of non-targeted metabolomics.

## Introduction

Plants, bacteria and animals emit complex mixtures of Volatile Organic Compounds (VOCs) forming blends or scent profiles. The chemodiversity of plant scent profiles is enormous as the last list of compounds published classifies over 1700 compounds (Knudsen et al., 2006). The variety of combinations in terms of quality and quantity of VOCs make many scent profiles unique for a species, or variety.

The structure of a scent profile is determined by a combination of three factors. First, developmental processes underlie the structure of a scent profile, as leaves, roots, flowers or fruits of a given plant emit distinct combinations of VOCs. Second, environmental conditions modify scent emission. For instance, some VOCs are typically produced under pathogen attacks (Kessler and Baldwin, 2002; Shimoda et al., 2012; Groen et al., 2016) and scent emission is affected by temperature and circadian regulation (Kolosova et al., 2001; Cna’ani et al., 2014; Terry et al., 2019a). Finally, genetic diversity plays a key role as many species, varieties and mutants emit differing scent profiles. Scent profiles can be used to identify species as it is a stable character and the major VOCs emitted tend to be a shortlist of metabolites (Knudsen et al., 2006; Raguso et al., 2006; Weiss et al., 2016a).

Whilst core scents are formed by a given blend of VOCs and are typical of a species or an organ, volatiles emitted in a non-constitutive way may play important biological roles. Thus, the combination of genetic diversity, morphogenesis and environmental cues, can make challenging the unequivocal determination of a scent profile emitted by a species, an organ, or under certain environmental conditions.

Reaching a consensus among samples of which compounds are comprising the constitutive metabolomic profile and which form the non-constitutive metabolome or discriminate between two sets of samples is mainly done manually. This causes two major problems, first, criteria are not always obvious or consistent and second, the procedure is extremely tedious, time consuming and prone to error. Sample size is key to determine accurately metabolic profiles. However, due to the difficulties found in data processing, sample size increment (n≥5) is neglected in many studies. An additional issue is the complexity of names given to a single chemical compound. In many cases, they include a common name, a chemical structure and sometimes isomers. The Chemical Abstract Service Number or CAS number is a single identifier that allows unambiguous assignation of a chemical structure. Thus the adoption of CAS-number defined metabolomes is the most appropriate way to produce metabolomics raw data in a suitable format for FAIR data management where data can be reanalysed (Wilkinson et al., 2016).

Here we provide an R-package that uses as inputs spreadsheet files produced by GC-MS apparatus to determine the core metabolome and non-constitutive compounds emitted by a set of samples. It compares between samples to give a set of common and differential set of metabolites in an automatic fashion.

We demonstrate the utility of gcProfileMakeR by analysing two genotypes of *Antirrhinum majus* affecting petal identity (Bey et al., 2004; Manchado-Rojo et al., 2012). Furthermore, we analysed the complete scent emission profile of *A.majus* lines with downregulated levels of *AmLHY* (Terry et al., 2019b). Our results indicate that the organ identity gene *deficiens* (Sommer et al., 1990), required to establish a petal organ identity, has a major impact in the scent profile emitted by the flowers, that result in scent profiles more similar to vegetative tissues. The downregulation of *AmLHY* causes among other features the appearance of volatiles undetected in the wild type plants, indicating a major coordination of scent emission by *AmLHY*. Using Machine Learning, we were able to classify the constitutive scent profiles of four genotypes with 0% error suggesting a great potential of gcProfileMakeR for downstream bioinformatics processing of metabolomic data.

## Results and Discussion

The full implementation of non-targeted metabolomics can give as a result a very large lists of liquid and/or gas chromatograms comprising hundreds of compounds (Zhu et al., 2018). Oftentimes, the number of compounds described undergo an arbitrary cut-off as major and minor components. The second reason to define only a subset of metabolites is that the comparison between samples is performed manually. We developed gcProfileMakeR, a tool accelerating the actual identification of common compounds in a set of samples. It uses reproducible criteria for downstream processing and data reusability. gcProfileMakeR was developed as an R package as R is open source, and the scientific community, especially biology, is doing a massive use of it. gcProfileMakeR determines the core metabolome and non-constitutive compounds present in a set of samples, thus allowing extensive exploration.

### gcProfileMakeR workflow

gcProfileMakeR uses two types of raw data: either XLS data files obtained directly from Agilent Chemstation software (Library Search Report) or CSV files (Fig. 1A). An example dataset can be retrieved within the library.

**Figure 1.**
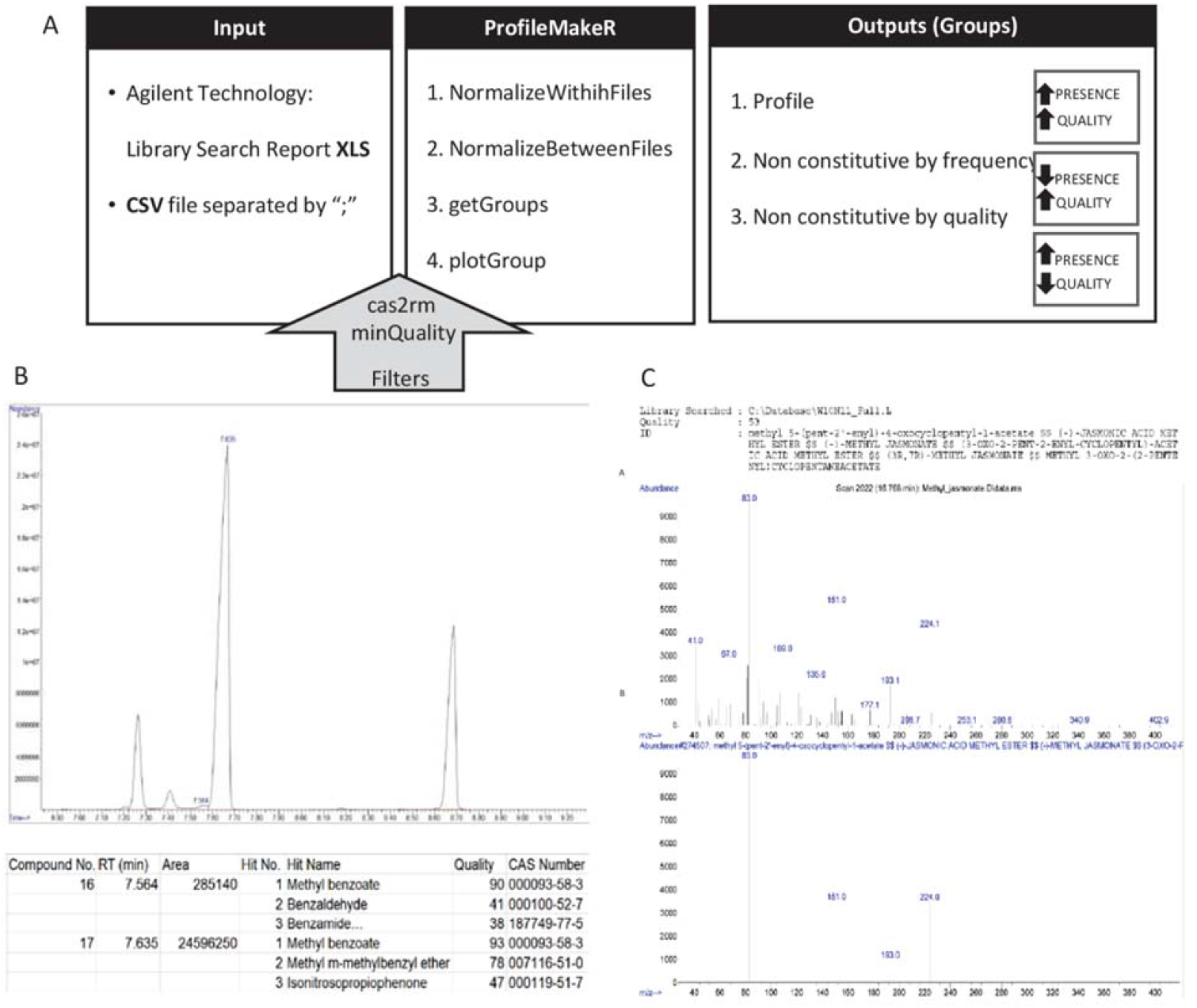
**(A)** gcProfileMakeR pipeiine. This library accepts Excel (.xls) and .csv files as input data. The first function, NormalizeWithinFiles, reads the data and groups compounds with similar retention time (RT) and common CAS numbers. Users also can apply two filters: cas2rm (compound/s to exclude) and minQuality (minimum quality). NormalizeBetweenFiles groups compounds with similar RT in all files, with the most representative CAS number. getGroups determines the constitutive and non-constitutive profiles (i.e. volatile profile) by frequency and quality, which are choose by the user. Finally, plotGroup graphs the constitutive, non-constitutive by frequency and/or non-constitutive profile by quality. (8) A standard chromatogram where two close peaks are integrated separately by default and dataset corresponding to peaks, where the identity with highest probability of the peaks is the same, methyl ben2oate (CAS number 93-58-3). (C) Mass spectra of methyl jasmonate(CAS No: 39924-52-2), a commercial standard (upper panel) and mass spectral database (lower panel)WilleylOth-NISTllb.

GC basic data contains information for each integrated peak about retention time (RT) and area of the peak. Mass spectra alignment with available libraries (MS libraries) allows to identify the compounds present in the sample with a certain degree of confidence (quality). Annotated compounds (hits) are listed according to the quality of the match between the mass spectra obtained and the mass spectra listed in the MS library. Hits are specified by chemical names of compounds and the CAS Registry Number associated to the hit/compound. CAS numbers are specific for a compound whereas chemical names are redundant and may imply different isomers or molecules. gcProfileMakeR works with RT, qualities and CAS numbers in order to provide lists of compounds identified by CAS numbers, areas and qualities. Chemical names are linked to the CAS numbers as they are understandable by scientists.

Two filters can be applied to pretreat data (Fig. 1A). The first one, cas2rm, will sort out any CAS number defined by the user, thus allowing the elimination of known contaminants, or compounds that are ubiquitous and complicate further analysis. The second filter, minQuality, eliminates hits, either first or secondary, with a quality below a defined level. It could leave retention times empty if being too strict (e.g. = 95). It allows to use a strategy of low strictness at the integration step and explore the data, decreasing the threshold to define a complete metabolome.

gcProfileMakeR uses three functions (Fig. 1A). The first function NormalizeWithinFiles, analyses each file/sample assigning for each retention time a set of possible hits (compounds). Peak areas of the same compounds with an identical CAS number found in different RTs, will be added (Fig. 1B). The second function NormalizeBetweenFiles, reaches a consensus between files in such a way that the same compounds separated in relatively close retention times are grouped together. The third function getGroups, establishes what is considered as “Profile”, “Non-constitutive by Frequency” and “Non-constitutive by Quality”. The rationale behind including a Non-constitutive by Quality list is that some compounds, even as chemical standards, give low quality due to poor representation in MS libraries, for instance methyl jasmonate (Fig. 1C). Frequency and quality default thresholds can be adjusted, thus allowing data exploration.

Default values have been tested with different sets of samples and number of samples and have proved the best outputs when compared to manual annotation (data not shown). The output of gcProfilemakeR are three mutually exclusive lists of compounds. The first set of compounds listed as “Profile” are those compounds which appear in all the samples of a given type i.e. genotype and/or treatment and which have a high matching quality: above a percentage of samples defined by the researcher. Compounds listed as “Non-constitutive by Frequency” are metabolites with a high mean-quality score (default: >85%) in the MS analysis but present in less than the percentage of the samples defined previously (Fig. 1A). Finally, compounds listed as “Non-constitutive by Quality” are metabolites with a low mean-quality (default: <85%) that are in at least 30% of the samples (default value). Frequency and quality thresholds can be adjusted for stringency thus allowing data exploration. Results can be plotted with the function plotGroup (Fig 2). In this function, compoundType parameter can be adjusted in order to get profiles (p), non-constitutive by frequency (ncf) or non-constitutive by quality (ncq). Results are plotted according to the average area and quality of each compound grouped in each category. The graphic obtained is in HTML format and allows, by pointing at the columns, to see the actual compound names that are linked to a CAS number (Fig. 2). Pointing at the quality percentages it shows the error rates of the quality for a given CAS number. This facilitates working with the graphics. They can also be saved as .png.

**Figure 2.**
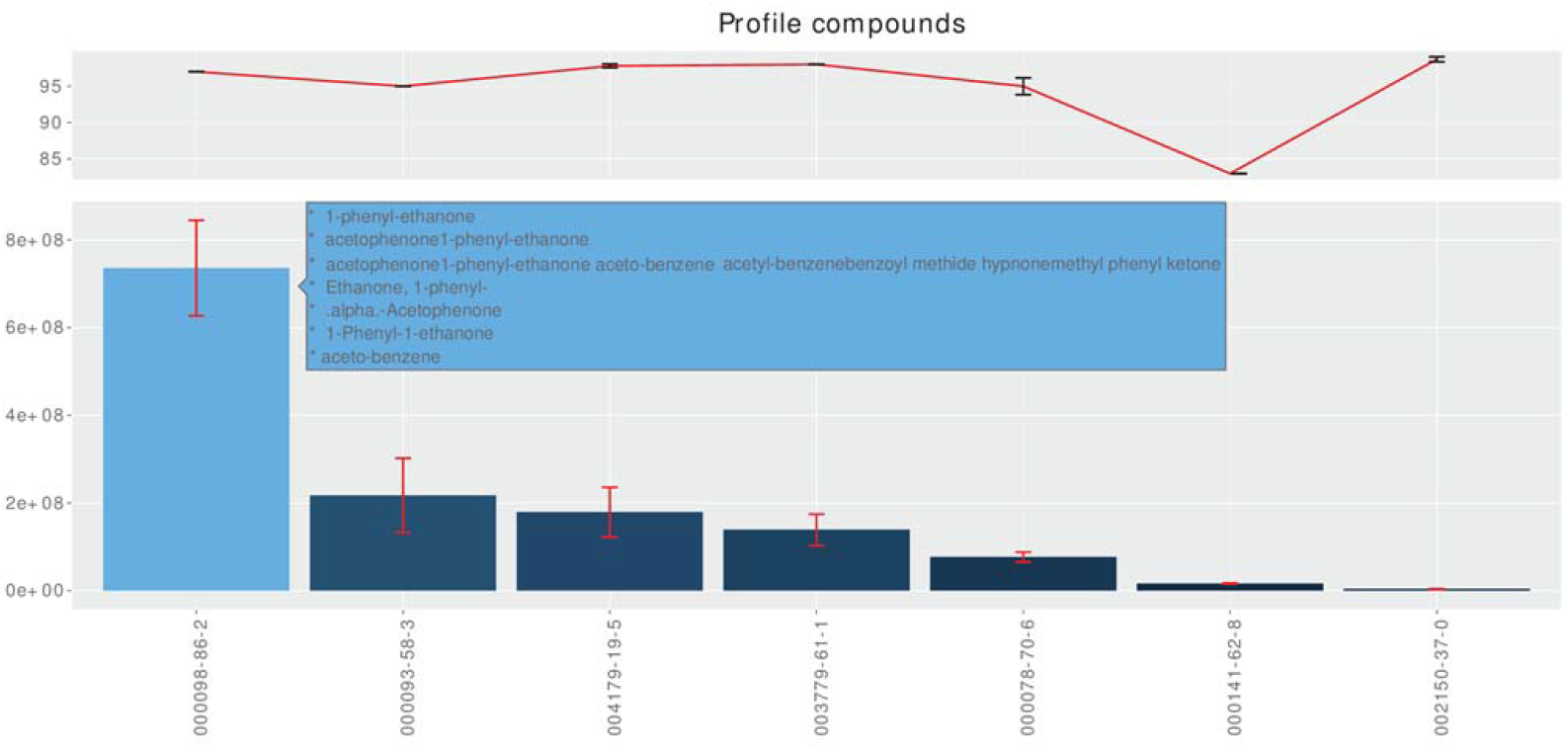
plotGroup function. This graph shows the constitutive profile by frequency of the wildtype snapdragon at ZT3 (*Zeitgeber* time). The x-axis shows the CAS number of volatile organic compounds. The upper part displays the average quality of volatiles (red line) and the lower part of the graph indicates the average areas of compounds (blue bars), that are plotted in decreasing order.

### Testing gcProfileMakeR in floral organ identity mutants and clock transgenic lines

We have experimentally validated gcProfileMakeR using a set of *Antirrhinum majus* mutants, transgenic and wild type plants. Floral scent emission depends on properly formed petal tissues, as weak alleles of B-function genes such as *deficiens-nicotianoides (def-nic)* (Sommer et al., 1991) or *compacta (co)*, show significant changes in the quantities of the terpenoids myrcene and ocimene, and the phenylpropanoid methyl benzoate (Manchado-Rojo et al., 2012). However, the complete scent profile had not been analysed.

We analyzed four datasets of floral volatiles, one corresponding to Sippe 50 wild types, one produced by the mutant *def-nic*, a third corresponding to *co* and a fourth corresponding to *RNAi:AmLHY*. We used a list of possible contaminants, which might proceed from the twister absorption matrix (Supplemental Table S1), and cas2rm to eliminate from our results any CAS numbers corresponding to siloxane or related derivatives.

Using gcProfileMakeR allowed to obtain a comprehensive profile present in at least 70 % of the samples (pFreqCutoff= 0.70). The wild type scent profile comprised seven constitutive VOCs in wild type flowers including benzenoids/phenylpropanoids and monoterpenes (Fig 3). In contrast, *co* produced only five, losing one benzenoid and one monoterpene, while it emitted decanal, an aldehyde absent in wild type. The stronger B-function mutant allele *def-nic* did not emit monoterpenes and had yet increased levels of aldehydes with presence of nonanal and decanal (Fig. 3). In sharp contrast, the scent profile of *RNAi:AmLHY* was significantly more complex than the wild type, and it included a total of fourteen VOCs comprising aldehydes, benzenoids/phenylpropanoids, mono and sesquiterpenes (Fig. 3).

**Figure 3.**
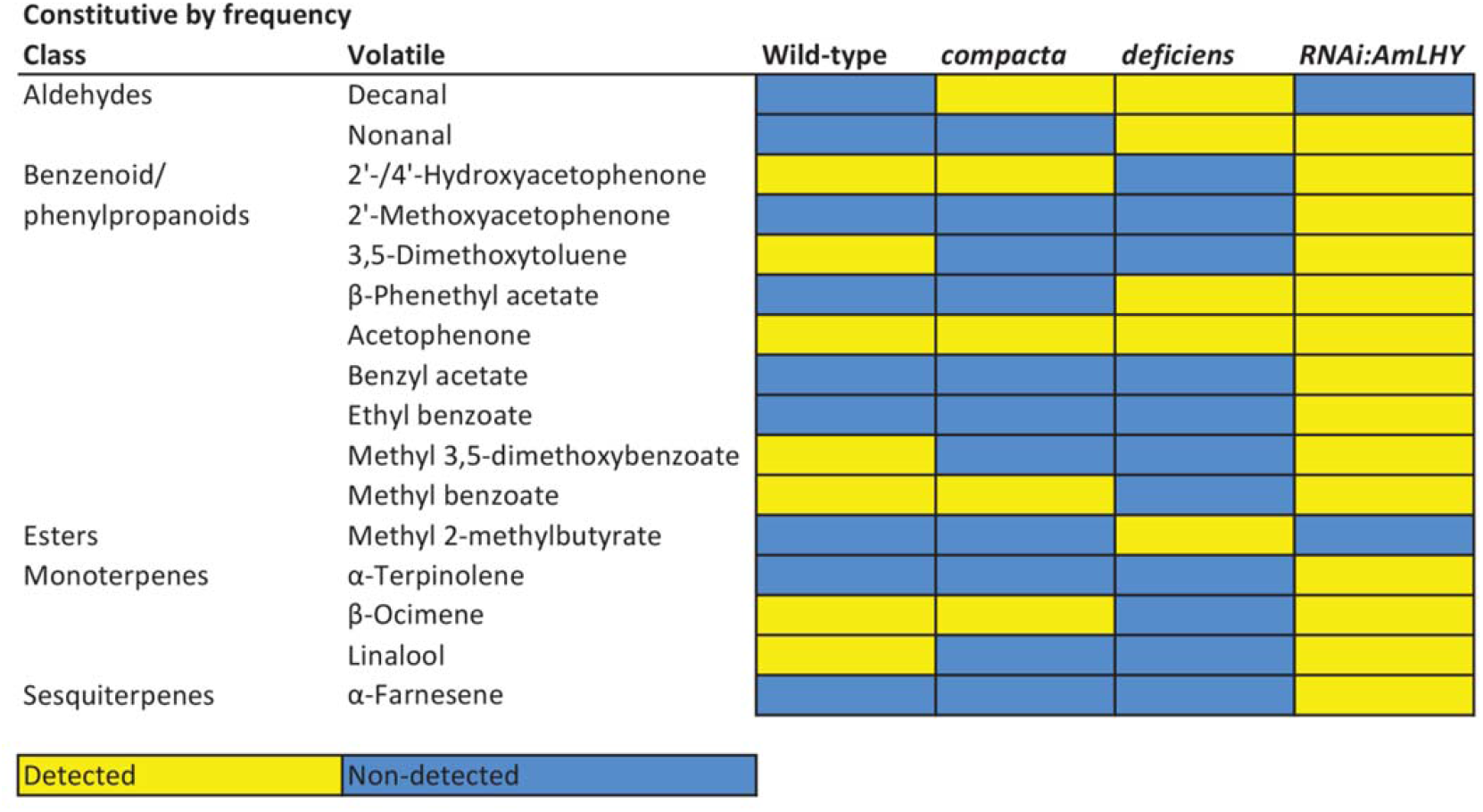
Heat map of constitutive by frequency scent profile of wild-type snapdragon (SIPPE5O), the mutants compacts and deficiens-nicotianoides and the transgenic line *RNAi:AmLHY. We* set minQuality to 80% (NormalizeWithinFiles function). Constitutive profile comprises those compounds that were present on at least the 70% of analyzed samples. Volatile compounds are clustered by class. Yellow and blue colors denote a detected and a non-detected compound, respectively.

When we inspected the Non-Constitutive by Frequency VOCs i.e. those found in less than 70% of the samples (Fig. 4), we found that wild type flowers emitted an additional set of five VOCs comprising the amine indole, benzenoids/phenylpropanoids and monoterpenes. In contrast, the number of volatiles emitted as Non-constitutive by Frequency by the rest of genotypes was substantially larger. The mutant *co* emitted 38 additional VOCs in all the categories including cycloalkanes such as cyclododecane, esters like borneol acetate, sesquiterpenes such as bornene and alpha-farnesene and terpene derivatives such as hexahydrofarnesyl acetone. The weak *def-nic* produced 19 volatiles while the *RNAi:AmLHY* lines produced 20 additional VOCs. Some of these were found only in the *RNAi:AmLHY* lines (see below). The analysis of non-constitutive by quality scent profile revealed 21 new volatiles (Supplemental Fig. S1). These compounds included aldehydes detected in transgenic lines, such as dodecanal and octanal.

**Figure 4.**
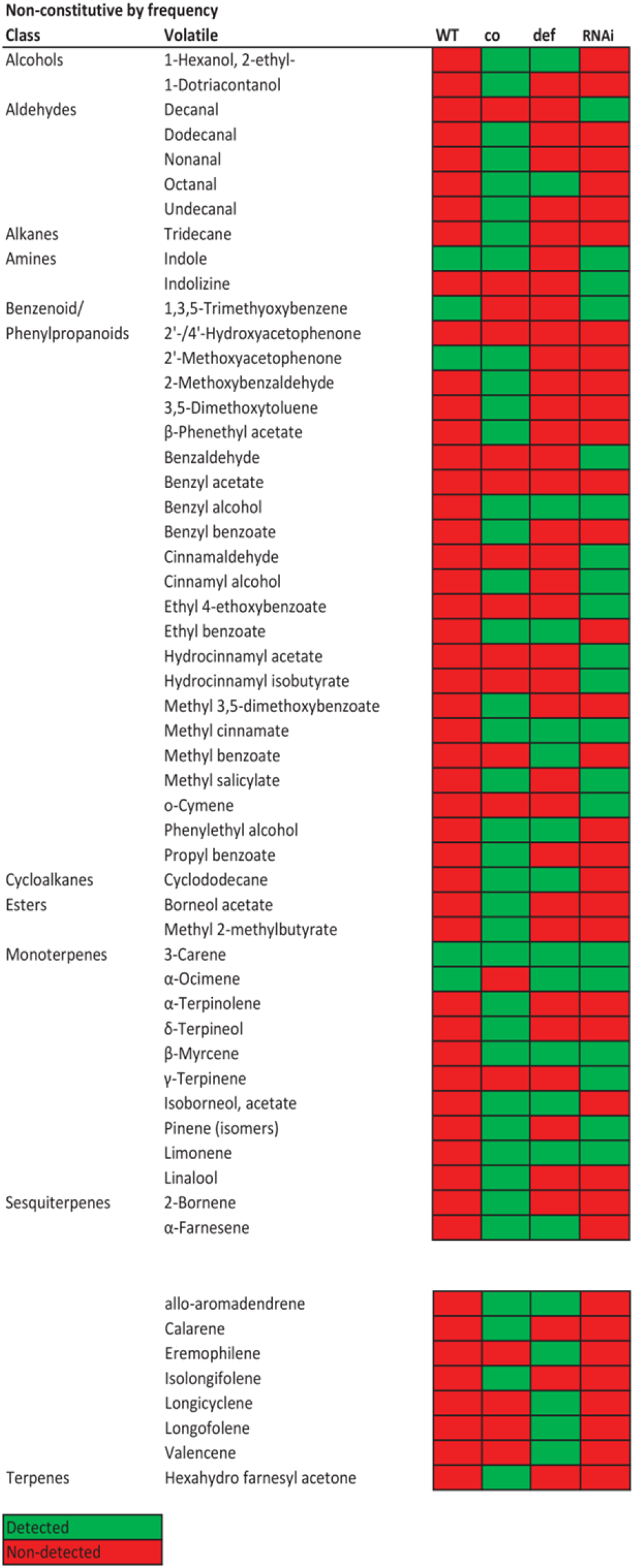
Heat map of non-constitutive by frequency scent profiles of wild-type snapdragon (Sippe SO, WT), the mutants co and def^nic^ and the transgenic line *RNAi:AmLHY (RNAi)*. We set minQualityto 80% (NormalizeWithinFiles function). Non-constitutive profile comprises those compounds that were present on or less than the 30% of analyzed samples. Volatile compounds are clustered by class. Green and red colors indicate a detected and a non-detected compound, respectively.

An important outcome of the data analyzed is that the actual genetic capacity of VOC emission in a wild type plant may be grossly underestimated. While the constitutive scent profile of the wild type is more complex than in the mutants, it is far simpler than the *RNAi:AmLHY* plants. This phenotype was also noticeable analyzing the daily emission of wild type and transgenic lines. The complexity of the constitutive and non-constitutive profile by frequency was higher in *RNAi:AmLHY* flowers (Fig. 5, Supplemental Fig. S2). Interestingly, we also found differences in time emission, as in case of the monoterpene linalool, that was not detected at ZT9 and ZT15 in wild type flowers but was constitutively emitted in all analyzed time points in transgenic snapdragons (Fig. 5).

**Figure 5.**
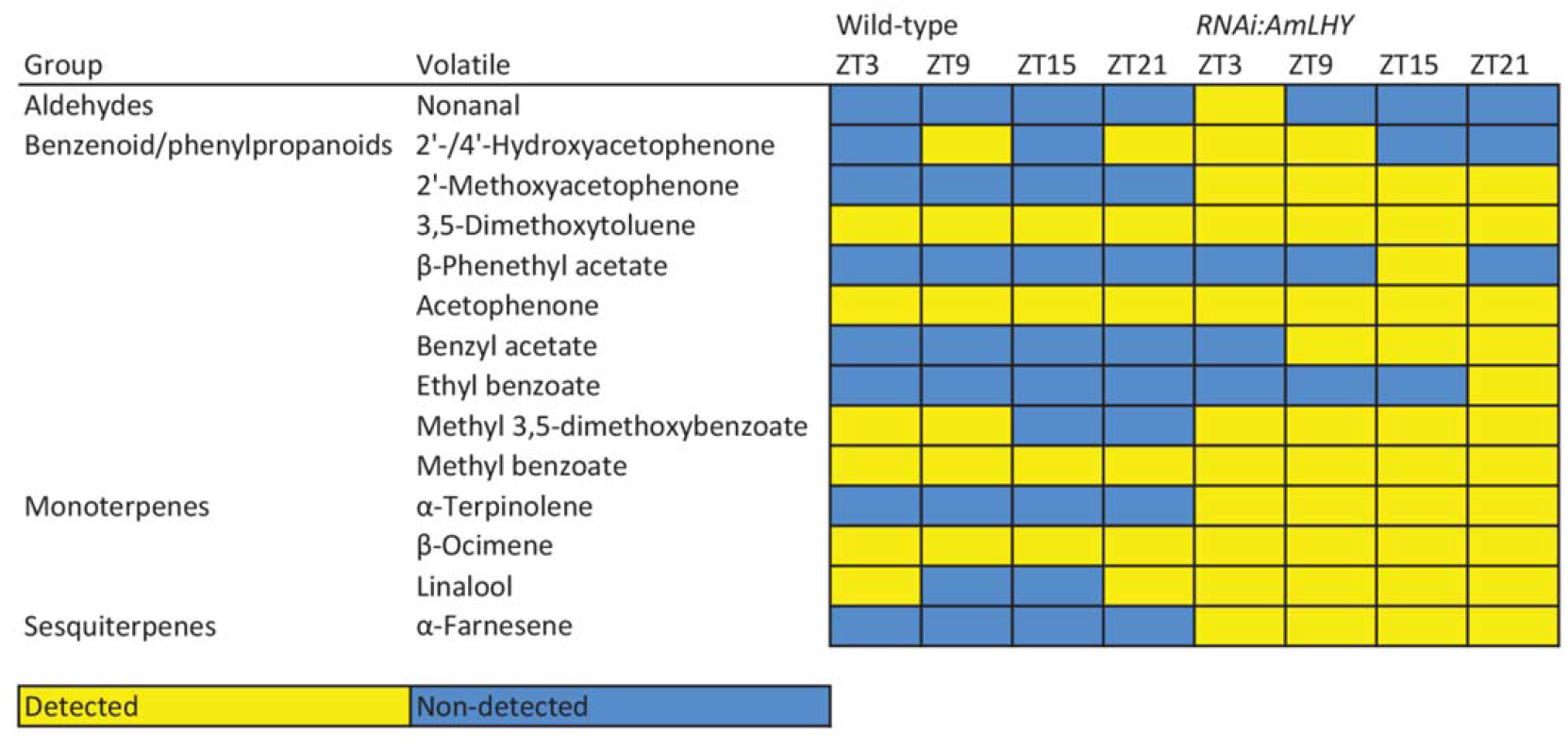
Constitutive scent profile of wild-type and transgenic *RNAi:AmLHY* snapdragons at four different time-points, denoted as ZT *(zeitgeber* time) 3, 9, 15 and 21. ZT0 represents the time of lights on and ZT12, lights off. We set minQuality to 80% (NormalizeWithinFiles function). Constitutive profile includes VOCs that were present on at least the 70% of analyzed samples. Volatiles are listed according to their class. Yellow indicates detected compounds and blue, non-detected compounds.

This suggests a general function of *DEF*, *CO* and *AmLHY* in establishing a concrete aroma typical of *A.majus* flowers.

### Effect of floral organ identity mutants and *RNAi:AmLHY* on VOC biosynthetic pathways

We identified compounds in snapdragon fragrance that are precursors of other volatiles, as benzaldehyde and its derivatives benzyl alcohol, benzyl acetate and methyl benzoate (Muhlemann et al., 2014). We found that, in general, snapdragon showed constitutive volatiles such as acetophenone, whereas other volatiles such as benzyl alcohol and methyl salycilate were present in the non-constitutive profile by frequency (Fig. 6). Based on previous data, we plotted the schematic pathway of benzenoid/phenylpropanoids and terpenoids pathways (Fig. 6, Fig. 7), indicating which group of snapdragon flowers emitted or not a compound, and its frequency among the analysed population. These results suggest a preferred route: the volatiles benzaldehyde and benzyl alcohol were not found in the constitutive profile of any snapdragon group whereas methyl benzoate was constitutively emitted in wild-type, compacta and *RNAi:AmLHY* lines but not in *deficiens-nicotianoides (def-nic)* group.

**Figure 6.**
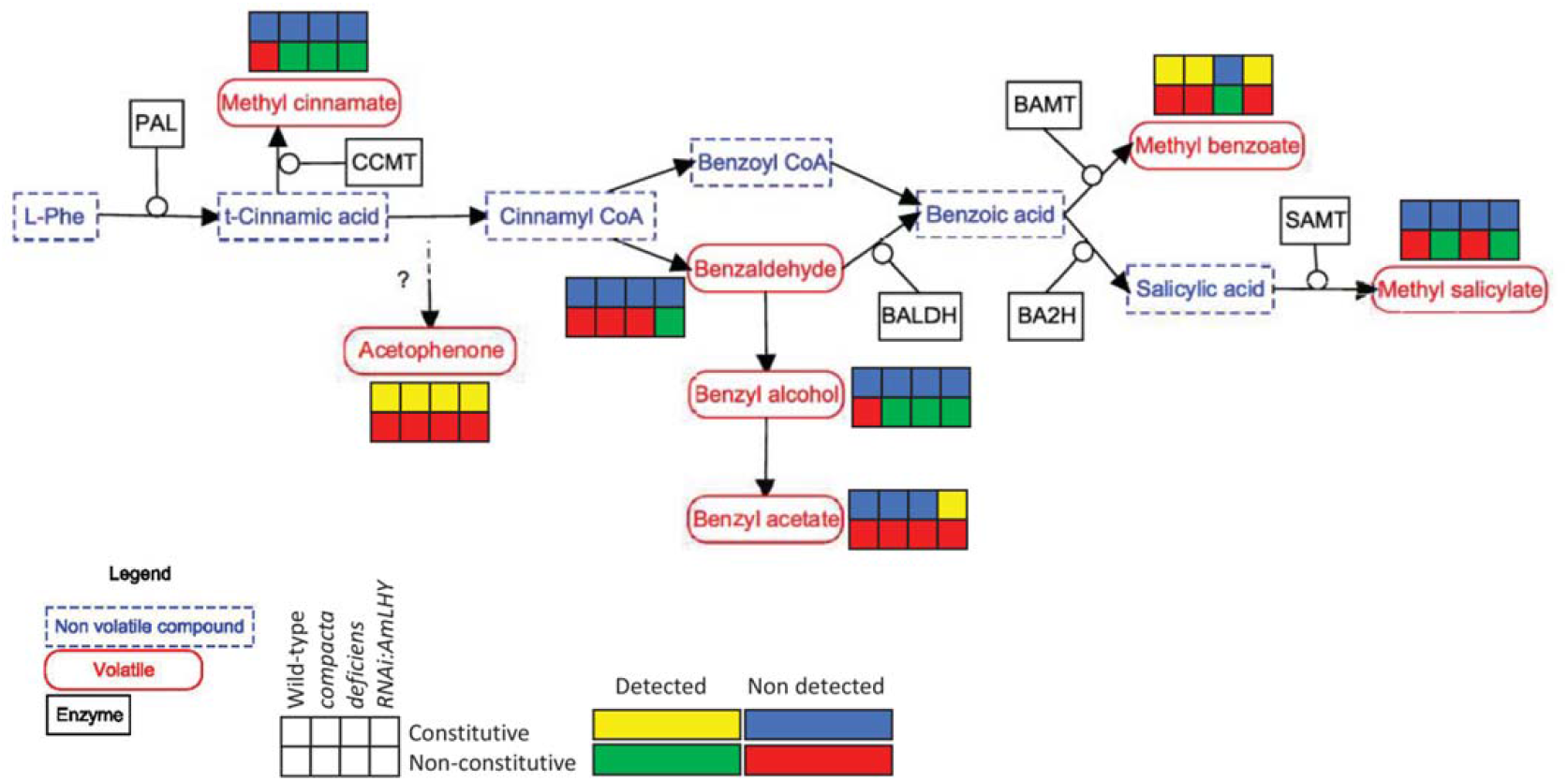
Benzenoid/phenylpropanoids schematic pathway. Detected and non-detected volatiles are shown as follow: first row refers to constitutive profiles and second row to non-constitutive by frequency profiles. Detected compounds in the constitutive and non-constitutive profiles are depicted by yellow and green, respectively. Non-detected compounds in the constitutive and non-constitutive profiles are indicated in blue and red, respectively. Each column represents a snapdragon group: wild-type (1st), *co* (2nd) and *def^nic^* (3rd) and transgenic lines *RNAi:AmLHY* (4th). PAL: phenylalanine ammonialyase, CCMT: cinnamic acid carboxyl methyl transferase, BALDH: benzaldehyde dehydrogenase, BA2H: benzoic acid 2-hydroxylase, BAMT: benzoic acid carboxyl methyl transferase, SAMT: salicylic acid carboxyl methyl transferase.

**Figure 7.**
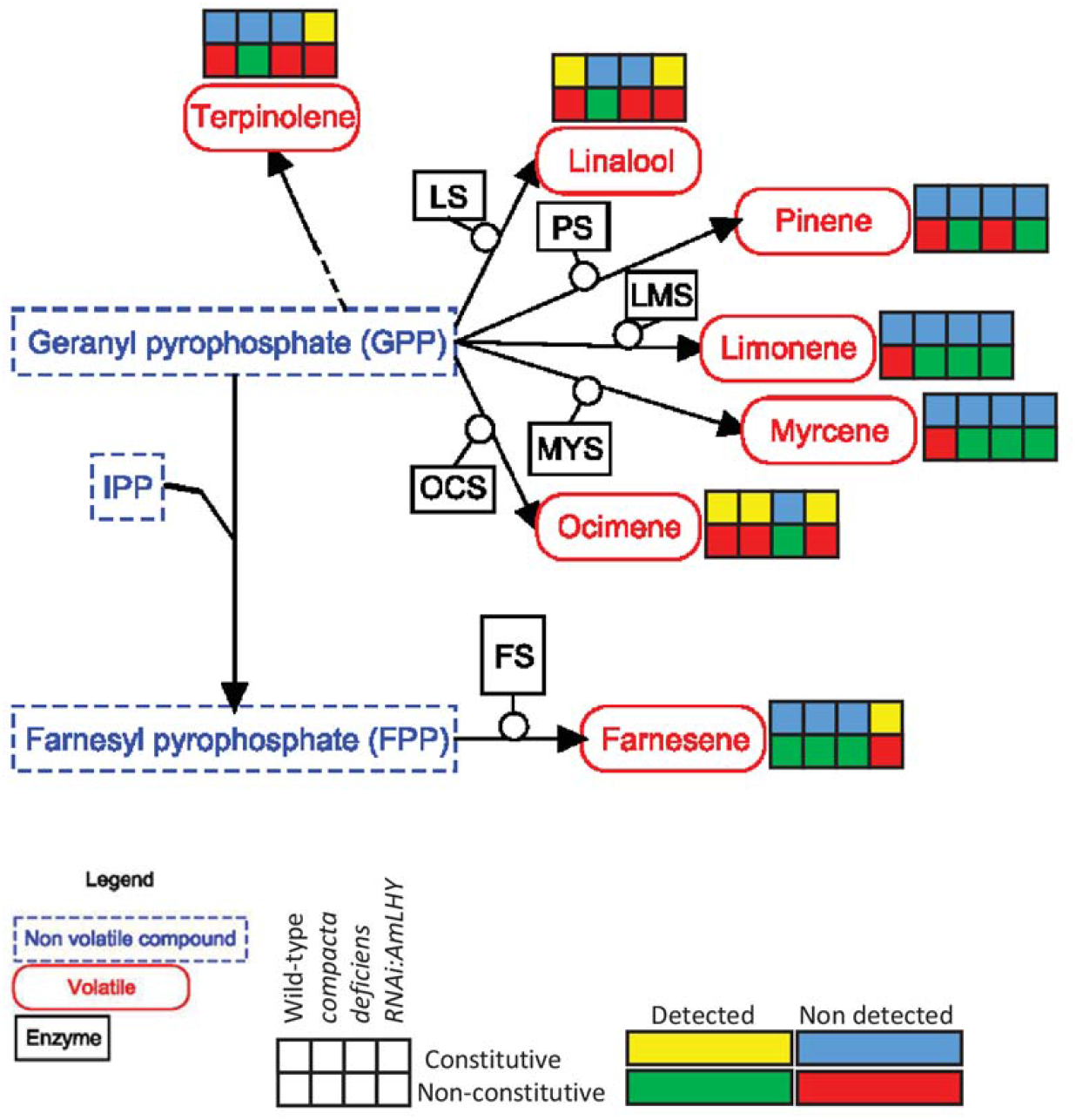
Terpenoids schematic pathway. Representations are like in Figure 6. LS: linalool synthase, PS: pinene synthase, LMS: limonene synthase, MYS: myrcene synthase, OCS: ocimene synthase, FS: farnesene synthase, IPP: isopentenyl diphosphate.

On the other hand, the monoterpenes linalool, pinene, limonene, myrcene and ocimene share the substrate geranyl pyrophosphate (Fig. 7). Pinene, limonene and myrcene were not present in the constitutive profile of analysed plant groups whereas linalool showed a constitutive emission in wild-type and *RNAi:AmLHY* and ocimene, in all plants except in *def-nic* mutant group. Differences in the constitutive and non-constitutive profile may be useful for further analysis of transcription factors, enzymes and transporters involved in volatile emission.

### Genotypes can be separated by Machine learning

Once we obtained a Constitutive Profile list of volatiles, we performed a classification analysis using the Machine Learning algorithm Random Forest (Breiman, 2001). Our data revealed that all snapdragon scent profiles were correctly classified (error out of bag or OOB, 0%) (Table 1). The “randomForest” package also provides a rank list with the accuracy in which a predictor, a volatile in our case, can be used for classification (Table 2). Altogether, our results show that gcProfileMakeR gives as output classified scent profiles that are sufficiently different to be separated by Machine Learning.

**Table 1.**
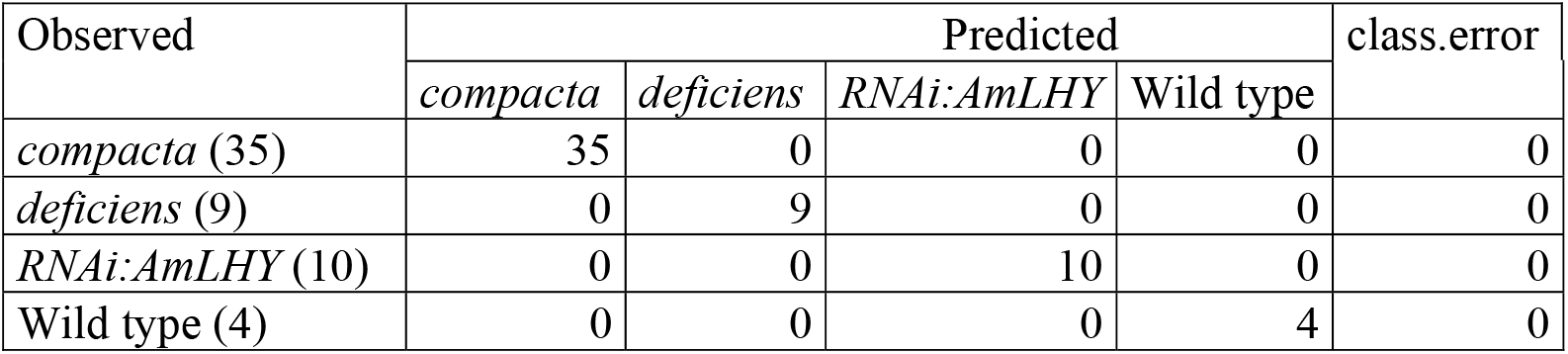
Random forest confusion matrix. The total number of samples of each snapdragon group is shown in parentheses (observed column). The number of misclassified samples of each group are in columns (predicted columns). The class.error column indicates the percentage of misclassified samples (1-[(total correct predictions/total predictions) x 100]).

| Observed | Predicted |  |  |  | class.error |
| --- | --- | --- | --- | --- | --- |
|  | *compacta* | *deficiens* | *RNAi:AmLHY* | Wild type |  |
| *compacta* (35) | 35 | 0 | 0 | 0 | 0 |
| *deficiens* (9) | 0 | 9 | 0 | 0 | 0 |
| *RNAi:AmLHY* (10) | 0 | 0 | 10 | 0 | 0 |
| Wild type (4) | 0 | 0 | 0 | 4 | 0 |

**Table 2.** Importance ranking of volatile organic compounds among *Antirrhinum majus* groups (wild-type, *compacta* mutant, *deficiens* mutant and *RNAi:AmLHY)* using random forest algorithm. The NIST library identifies two pairs of similar compounds which share the same retention time, 2’-Hydroxyacetophenone and 4’-Hydroxyacetophenone, and 2’-Methoxyacetophenone and 4’-Methoxyacetophenone, respectively. These compounds are depicted with a slash (“/”) in the table. Volatiles are ranked based on mean decrease in accuracy (MDA). This value indicates the accuracy in which a volatile can be used for classification.

| Volatile | MDA |
| --- | --- |
| Nonanal | 16.26 |
| Farnesene | 14.8 |
| Methyl 2-methylbutyrate | 14.65 |
| 3,5-Dimethoxytoluene | 14.44 |
| Methyl benzoate | 12.66 |
| Acetophenone | 10.96 |
| Phenethyl acetate | 9.57 |
| Methyl 3,5-dimethoxybenzoate | 9.26 |
| Ocimene | 6.52 |
| Linalool | 5.99 |
| Decanal | 5.54 |
| 2'-/4'-Hydroxyacetophenone | 5.04 |
| 2'-/4'-Methoxyacetophenone | 4.93 |
| Terpinolene | 3.75 |
| Ethyl benzoate | 1.42 |
| Benzyl acetate | 0 |

As sessile organisms, plants rely on their chemistry to deal with the many interactions conditioning their survival (abiotic and biotic). That may be the reason why plant volatile chemotypes are known for their variability with regard to composition and relative abundance of VOCs (Junker et al., 2018). gcProfileMakeR should ease the task to define constitutive and non-constitutive metabolomes in large datasets.

## Materials and Methods

### Plant material and VOCs analysis

We used flowers from *Antirrhinum majus compacta (co)* and *deficiens-nicotianoides (def-nic)* mutants (Manchado-Rojo et al., 2012) and *RNAi:AmLHY* from three independent transgenic lines (Terry et al., 2019b). *Antirrhinum* plants were grown in the greenhouse as described previously, using standard methods (Weiss et al., 2016b). Scent samples were analysed according to (Ruiz-Hernández et al., 2017). Samplings periods of VOCs were 24 hours for *def-nic* and *co*, while WT and *RNAi*: *AmLHY* were sampled every six hours for a complete day. The *RNAi:AmLHY* samples were aggregated to compare to other genotypes. We analysed 16 biological replicas for wild type Sippe 50, 35 for *co*, 9 for *def-nic* and 40 for *iRNA:AmLHY*.

### gcProfileMakeR

gcProfileMakeR R package is available at git clone git@gitlab.atica.um.es:fernando.perez8/gcProfileMakeR.git.

Some packages are recommended to be pre-installed in R before gcProfileMakeR runs: readxl, plyr, stringr, dplyr, tidyr, ggplot2 and egg.

### Machine Learning Analysis

We used the random forest algorithm implemented in the R package “randomForest”” (Liaw and Wiener, 2002) (R version 3.6.1). We rearranged our data in a data frame where the rows correspond to the samples from wild type, *co*, *defnic* mutants and *RNAi:AmLHY* and the columns contain the value of the selected volatiles, expressed as integrated peak area divided by the fresh weight. We used *randomForest* default parameters, setting parameter “importance” as “TRUE” and obtaining a classification random forest.

## Notes

Funding information: This work was supported by the Ministerio de Economía Industria y Competitividad [BFU2017-88300-C2-1R] to [MEC] [JW], [BFU2017-88300-C2-2R] to [PJN]; Fundación Séneca [19398/PI/14] to [MEC] and PhD grant by the Ministerio de Educación Cultura y Deporte [FPU13/03606] to [VRH].

## Parsed Citations

Bey M, Stuber K, Fellenberg K, Schwarz-Sommera Z, Sommer H, Saedler H, Zachgo S (2004) Characterization of Antirrhinum petal development and identification of target genes of the class B MADS box gene DEFICIENS. Plant Cell 16: 3197–3215

Breiman L (2001) Random Forests. Machine Learning 45: 5–32

Cna’ani A Mühlemann JK, Ravid J, Masci T, Klempien A, Nguyen TTH, Dudareva N, Pichersky E, Vainstein A (2014) Petunia x hybrida floral scent production is negatively affected by high-temperature growth conditions. Plant, cell & environment. doi: 10.1111/pce.12486

Groen SC, Jiang S, Murphy AM, Cunniffe NJ, Westwood JH, Davey MP, Bruce TJA, Caulfield JC, Furzer OJ, Reed A, et al (2016) Virus Infection of Plants Alters Pollinator Preference: A Payback for Susceptible Hosts? PLOS Pathogens 12: e1005790

Junker RR, Kuppler J, Amo L, Blande JD, Borges RM, Dam NM van, Dicke M, Dötterl S, Ehlers BK, Etl F, et al (2018) Covariation and phenotypic integration in chemical communication displays: biosynthetic constraints and eco-evolutionary implications. New Phytologist 220: 739–749

Kessler A, Baldwin IT (2002) Plant responses to insect herbivory: the emerging molecular analysis. Annual Review of Plant Biology 53: 299–328

Knudsen JT, Eriksson R, Gershenzon J, Ståhl B, Stahl B (2006) Diversity and distribution of floral scent. Botanical Review 72: 1–120

Kolosova N, Gorenstein N, Kish CM, Dudareva N (2001) Regulation of circadian methyl benzoate emission in diurnally and nocturnally emitting plants. Plant Cell 13: 2333–2347

Liaw A, Wiener M (2002) Classification and regression by random Forest. R news 2: 18–22

Manchado-Rojo M, Delgado-Benarroch L, Roca MJ, Weiss J, Egea-Cortines M (2012) Quantitative levels of Deficiens and Globosa during late petal development show a complex transcriptional network topology of B function. The Plant journal:for cell and molecular biology 72: 294–307

Muhlemann JK, Klempien A, Dudareva N (2014) Floral volatiles: from biosynthesis to function. Plant, cell & environment 37: 1936–49

Raguso RA, Schlumpberger BO, Kaczorowski RL, Holtsford TP (2006) Phylogenetic fragrance patterns in Nicotiana sections Alatae and Suaveolentes. Phytochemistry 67: 1931–1942

Ruiz-Hernández Victoria, Hermans Benjamin, Weiss Julia, Egea-Cortines Marcos (2017) Genetic analysis of natural variation in Antirrhinum scent profiles identifies BENZOIC ACID CARBOXYMETHYL TRANSFERASE as the major locus controlling methyl benzoate synthesis. Frontiers in plant science 8: 27–40

Shimoda T, Nishihara M, Ozawa R, Takabayashi J, Arimura GI (2012) The effect of genetically enriched (E)-β-ocimene and the role of floral scent in the attraction of the predatory mite Phytoseiulus persimilis to spider mite-induced volatile blends of torenia. New Phytologist 193: 1009–1021

Sommer H, Beltran JP, Huijser P, Pape H, Lonnig WE, Saedler H, Schwarz-Sommer Z, Schwarzsommer Z, Beltrán JP, Huijser P, et al (1990) Deficiens, a Homeotic Gene Involved in the Control of Flower Morphogenesis in Antirrhinum-Majus - the Protein Shows Homology to Transcription Factors. EMBO Journal 9: 605–613

Sommer H, Nacken W, Beltran P, Huijser P, Pape H, Hansen G, Flor P, Saedler H, Schwarz-Sommer Z, Hansen R, et al (1991) Properties of Deficiens, a Homeotic Gene Involved in the Control of Flower Morphogenesis in Antirrhinum-Majus. Development Supplement 1: 169–175

Terry MI, Pérez-Sanz F, Díaz-Galián MV, Pérez de los Cobos F, Navarro PJ, Egea-Cortines M, Weiss J (2019a) The Petunia CHANEL Gene is a ZEITLUPE Ortholog Coordinating Growth and Scent Profiles. Cells 8: 343

Terry MI, Pérez-Sanz F, Navarro PJ, Weiss J, Egea-Cortines M (2019b) The Snapdragon LATE ELONGATED HYPOCOTYL Plays A Dual Role in Activating Floral Growth and Scent Emission. Cells 8: 920

Weiss J, Mühlemann JK, Ruiz-Hernández V, Dudareva N, Egea-Cortines M (2016a) Phenotypic space and variation of floral scent profiles during late flower development in Antirrhinum. Frontiers in plant science 7: 1903

Weiss Julia, Alcantud-Rodriguez Raquel, Toksöz Tugba, Egea-Cortines M (2016b) Meristem maintenance, auxin, jasmonic and abscisic acid pathways as a mechanism for phenotypic plasticity in Antirrhinum majus. Scientific reports 6: 2–11

Wilkinson MD, Dumontier M, Aalbersberg IjJ, Appleton G, Axton M, Baak A, Blomberg N, Boiten J-W, da Silva Santos LB, Bourne PE, et al (2016) The FAIR Guiding Principles for scientific data management and stewardship. Scientific Data 3: 160018

Zhu G, Wang S, Huang Z, Zhang S, Liao Q, Zhang C, Lin T, Qin M, Peng M, Yang C, et al (2018) Rewiring of the Fruit Metabolome in Tomato Breeding. Cell 172: 249–261.e12

